# Dual roles of electrostatic-steering and conformational dynamics in the binding of calcineurin’s intrinsically-disordered recognition domain to calmodulin

**DOI:** 10.1101/277236

**Authors:** Bin Sun, Eric C. Cook, Trevor P. Creamer, Peter M. Kekenes-Huskey

## Abstract

calcineurin (CaN) is a serine/threonine phosphatase that regulates a variety of physiological and pathophysiological processes in mammalian tissue. The CaN regulatory domain (RD) is responsible for regulating the enzyme’s phosphatase activity, and is believed to be highly-disordered when inhibiting CaN, but undergoes a disorderto-order transition upon diffusion-limited binding with the regulatory protein calmodulin (CaM). The prevalence of polar and charged amino acids in the regulatory domain (RD) suggests electrostatic interactions are involved in mediating CaM binding, yet the lack of atomistic-resolution data for the bound complex has stymied efforts to probe how the RD sequence controls its conformational ensemble and long-range attractions contribute to target protein binding. In the present study, we investigated via computational modeling the extent to which electrostatics and structural disorder co-facilitate or hinder CaM/CaN association kinetics. Specifically, we examined several RD constructs that contain the CaM binding region (CAMBR) to characterize the roles of electrostatics versus conformational diversity in controlling diffusion-limited association rates, via microsecond-scale molecular dynamics (MD) and Brownian dynamic (BD) simulations. Our results indicate that the RD amino acid composition and sequence length influence both the dynamic availability of conformations amenable to CaM binding, as well as long-range electrostatic interactions to steer association. These findings provide intriguing insight into the interplay between conformational diversity and electrostatically-driven protein-protein association involving CaN, which are likely to extend to wide-ranging diffusion-limited processes regulated by intrinsically-disordered proteins.

## 1. Introduction

calcineurin (CaN) is a ubiquitously expressed protein that regulates myriad developmental and signaling processes (1, 2). It is chiefly regulated by calmodulin (CaM), one of the most prolific enzymes in terms of its role in shaping intracellular signal transduction cascades. Despite the fundamental importance of CaM-regulated CaN phosphatase activity in organism physiology, the molecular mechanisms governing this process are incompletely understood. Given that CaM/CaN is a prototypical example of a protein/protein complex involving a globular protein (CaM) and an intrinsically disordered binding domain (CaN) (3, 4), structural details of the protein/protein complex are restricted to intact CaM bound to a small fragment of the CaN regulatory peptide. In this regard, the CaM/CaN complex is similar to the tens of CaM/protein target complexes (5) that have resisted structure determination methods beyond the binding of short peptides. Further, despite the CaN regulatory domain presenting little stabilized secondary structure, the CaM/CaN complex has a remarkably strong protein/protein affinity in the picomolar range (6), and paradoxically, the complex formation occurs rapidly in a diffusion-limited regime. In this paper, we utilize molecular simulations to probe how intrinsically disordered peptide (IDP) fragments from the CaN regulatory domain can achieve diffusion-limited binding kinetics to CaM. Our findings provide insight into a fundamentally important signal transduction pathway (7, 8), and specifically, how properties of an intrinsically disordered peptide (IDP) conformation ensemble and its ability to binding globular protein partners depend on the charge composition and electrostatic environment of the IDP. We anticipate these findings will be of broad importance to CaM-complexes involving IDP binding partners.

CaN is heterodimeric protein consisting of two domains: chain A (57-61 kDa) and chain B (19 kDa) (2, 7), while CaM (17 kDa) is comprised of two alpha-helix rich domains capable of binding Ca^2+^. At Ca^2+^ concentrations typical of resting cells (50 to 100 nM) (9), CaN phosphatase activity is negligible, while CaM is believed to be in Ca^2+^-free state (10). Under these conditions, the CaN catalytic domain is autoinhibited by the protein’s auto-inhibitory domain (AID). At rising Ca^2+^ concentrations, the CaN AID is removed from the catalytic domain. It is believed that binding of CaM to the AID-containing CaN (RD) (Ser373 to Thr468) is a critical determinant of this process (4, 11). However, the RD’s intrinsically disordered structure (3, 4), that is, it’s absence of significant folded structure (12, 13) has stymied efforts to probe its regulation by CaM. Similar to other IDP-containing complexes, well-defined secondary structure is presented only in the presence of a binding partner (12, 14–16).

It is becoming increasingly clear that the sequence composition of IDPs plays a profound role in defining the ensemble of conformations in equilibrium. In absence of hydrophobic residues (17) that would otherwise promote collapse of protein into a molten apolar core, many IDPs such as the CaN RD are polyampholytic (18). Such polyampholytic sequences feature abundant positive and negative-charged residues that favor solute/solvent interactions and thereby prevent the formation of folded structures. Recently, observations of high charge density in IDPs have culminated in a predictive metric, net charge per residue (NCPR), for relating attributes of IDP structure, such as compactness and shape, to sequence composition (19, 20). Formally, *NCPR* = |*f*+ −*f*−| where *f*+ and *f*− are fractions of positively and negatively charged residues, respectively, and fraction of charged residues (FCR) is calculated as 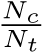 where *N*_*c*_ is the number of charged residues and *Nt* is the total number of residues. Generally, IDPs with large net charge per residue (NCPR) values (*>* 0.25) tend to adopt more extended conformations due to repulsive intra-molecular electrostatic interactions (19, 21). For CaN, the NCPR metric for the CaM binding region indicates that the RD subsection is in the “Janus” region (as shown in Fig. S1(d)) between collapsed and extended. It is important to note that the distribution of charged residues in the CaN RD is by no means uniform, thus local variations in the charge density along the RD sequence will likely determine the balance between extended and collapsed configurations. We postulated therefore that the NCPR near the RD CaM binding region (CAMBR) critically control the availability of “open” conformations amenable to CaM binding.

#### Significance Statement

Signal transduction pathways are frequently tuned through protein-protein interactions involving intrinsically disordered peptide (IDP) s. While quantitative insights into these important interactions are beginning to emerge, there remain significant unanswered questions, such as how intrinsically-disordered sequences can achieve diffusion-limited binding kinetics with folded partners, despite the paradoxical lack of structure resembling the bound complex. To answer this question, we utilized multi-scaled simulations technologies to reveal the role of electrostatics in driving diffusion limited association between intrinsically disordered region of calcineurin (CaN) with the ubiquitous calmodulin (CaM) protein. We found that the association kinetics between CaN and CaM are strongly controlled by long-range electrostatic interactions traditionally implicated in the rapid association of globular proteins, in addition to ‘local’ electrostatic interactions that tune access to IDP binding sites.

Protein-protein recognition events that fall within a diffusion-limited regime (22–25) commonly exhibit relatively minor structural changes as isolated proteins form the bound complex. Hence, it is surprising that several protein-protein interactions involving IDPs like the CaN RD form with diffusion-limited rates (26–28). Shammas et al (26) for instance have attributed the remarkably high basal association rate 2.1 *×* 10^7^ M^−1^ s^−1^ for a protein-protein interaction (PPI) involving an IDP partner to a loosely-associated ‘encounter-complex’ between the IDP and its target. It has been speculated that, in the unbound state, IDPs may adopt so called “residual structure” resembling the bound state (29) that provide the basis for rapid binding via conformation selection (30). Similarly, we recently determined via stopped-flow experiments(arXiv:1611.04080v1) for CaN binding to CaM that association is both diffusion-limited and dependent on ionic strength, as is typical of electrostatically-driven encounter complexes observed for PPIs from globular proteins (23, 31, 32). In contrast, other examples such as p53 up-regulated modulator of apoptosis (PUMA) do not adhere to this paradigm, as its rapid association rate depends on temperature and solvent viscosity in manners atypical of diffusion-limited association events (33–35). For PUMA, it was speculated that association is driven by an induced-fit mechanism that depends on tight interactions between binding partners (34), in contrast to the encounter complex suggested by Shammas et al. Hence, there is likely a system-dependent (36) interplay between properties of the IDP conformational ensemble, conformational dynamics and their diffusion-mediated encounters with binding partners that dictate observed PPI association rates.

In our recent study(arXiv:1611.04080v1), we implicated long-range, ionic strength-dependent electrostatic interactions as a basis for rapid association between rigid CaM and CaN peptide constructs. Based on our findings, we hypothesize here that RD sequence charge composition (as measured by NCPR) and ionic strength influence the dynamic availability of conformations amenable to CaM binding, while long-range electrostatic interactions drive diffusion-limited association (see Fig. 1). To investigate this hypothesis, we utilize long-timescale molecular dynamics (MD) simulations to probe the highly dynamic conformational ensembles comprising the RD constructs, toward delineating the extent to which conformational gating kinetics and long-range electrostatic interactions govern IDP/protein association. A chief outcome of this work is that we identify a quantitative link between ionic strength-dependent interconversion kinetics of the RD IDPs’ conformational ensemble with long-range PPI association. Overall, our data indicate that the RD IDP charge density influences the availability of CaM compatible configurations, that interconversion kinetics and thus presentation of the CaM-binding motif are rapid, and that long-range electrostatic interactions are instrumental for driving diffusion-limited association. As a consequence, we have clarified roles of charge density in CaM/CaN PPI kinetics, which may generalize to a variety of protein-protein interactions controlled by intrinsically-disordered binding partners.

**Fig. 1.**
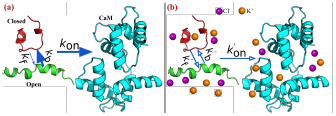
(a) Association rate between CaN peptide and CaM is determined by two components: the gating kinetics between open and closed state of CaN peptide (depicted by *k*_b_ and *k*_f_) and the diffusional encounter rate between open sate CaN peptide and CaM (depicted by *kon*). b) In the present study, we explore the role of electrostatic (tuned by varying ionic strengths) in modulating gating kinetics and diffusional encounter rate.

## 2. Results and Discussion

### A. Molecular simulations confirm the intrinsically-disordered structure of the CaN regulatory domain

Several studies using Fourier transform infrared spectroscopy, hydrolysis experiments and X-ray crystallography indicate that the nearly one-hundred amino acids of the CaN RD domain (Ser373 to Thr468 (3, 4)) form an intrinsically disordered ensemble (4, 37–39). The exception is a short stretch of roughly twenty amino acids (Ala391 to Arg414) comprising the CAMBR, which adopt an alpha-helix in the presence of CaM (40). Our study focuses on three RD constructs (pCaN, lpCaN and lpcCaN, see Fig. 2) that we have recently experimentally demonstrated assume diffusion-limited association with CaM (arXiv:1611.04080v1). Bioinformatics tools DisEMBL (41) and CIDER (42) suggest that like the intact RD domain, the RD constructs are likely to be disordered (Fig. S1). Moreover, as shown in Fig. 2, the constructs have similar FCR values (pCaN: 0.375, lpCaN: 0.382, lpcCaN: 0.382), but vary with regard to their charge compositions. For instance, pCaN and lpcCaN have similar NCPR values of 0.291 and 0.264, which are considerably larger than the value for lpCaN (0.088). Based on prior works (19, 21), NCPR scores above 0.25 are suggestive of extended IDP conformations given the propensity for repulsive intramolecular interactions, whereas those below this threshold are comparatively compact. We expected therefore that 1) the CaN peptides lack well-resolved secondary structure characteristic of a folded protein and 2) the ensemble of lpCaN should be modestly more compact than that of lpcCaN, given that latter has higher charge density.

To investigate the hypothesis, we performed 5 *μ*s MD simulations in triplicate (total 15 *μ*s) at 0.15 M and 1.5 M ionic strength, respectively. The choice of physiological (0.15 M) and high ionic strength was intended to probe the contribution of intra-peptide electrostatic interactions to ensemble properties, as such interactions would be screened at 1.5 M ionic strength. While simulations of IDPs of up to 100 residues have been reported elsewhere (21), the breadth of simulations used in this study restricted our construct sizes to 24 to 34 a.a. Our MD simulations indicate that the heavy atom root mean squared fluctuations (RMSF) for each residue in Fig. S1(e-f) are shown to be larger than 5 Å for all three peptides at both ionic strengths, which is consistent with the high mobility loop scores reported in Fig. S1(b). These data suggest that the peptides do not form stable folded structures in solution regardless of ionic strength.

**Fig. 2.**
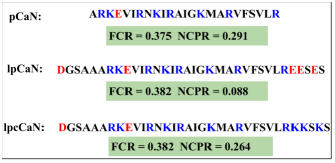
Amino acid sequences of three CaN peptide constructs studied in present study. pCaN: native CAMBR of CaN which spans Ala391 to Arg414. lpCaN: five residues affixed to the N/C-terminis of pCaN. lpCaN: mutant of lpCaN with three glutamic acids mutated to three lysines at the C-termini. The FCR and NCPR of each peptide were also shown‥

The MD-generated structures present a multitude of conformations, ranging from loosely-formed, hairpin-like configurations to extended structures. lpCaN presents perhaps the most hairpin character, as corroborated by intramolecular contacts reported through contact map analyses in Fig. S2. Among these contacts are prominent interactions between Arg12-Glu30, Arg23-Glu30 and Arg13-Glu32, which we attribute to transient salt-bridge formation. For lpcCaN, the mutation of negative residues (Glu30, Glu31 and Glu33) to the positively charged residues (Lys30, Lys31 and Lys33) appears to disrupt these intramolecular contacts, thereby yielding a more extended conformation ensemble relative to lpCaN. Given the similar NCPR values of pCaN and lpcCaN, we expected pCaN would similarly present fewer intramolecular contacts than lpCaN. Surprisingly, pCaN has similar contact map features as lpCaN, that is, both peptides have comparable intra-contacts. Later we will demonstrate that lpCaN and pCaN differ in terms of the kinetics of conversion between ensemble conformations. Additionally, we found that increasing ionic strength to 1.5 M screens the electrostatic interactions between residues comprising the reported salt bridges. As a result, we observe for pCaN and lpCaN that the structures become modestly more extended on average.

To support the formation of the CaN/CaM PPI, the CaN CAMBR must be revealed to the solvent-exposed CaM surface. The exposure of the CAMBR could occur spontaneously, which would promote binding by presenting mutually compatible conformations independent of the complementary species, or only in the presence of the binding partner CaM, which is more typical of an induced fit mechanism. In the previous section, we indicated that the peptides have considerable structural variability, therefore here we determine whether this variability confers greater access to the CAMBR *independent* of CaM.

In Fig. 3(a-c), we report the root mean squared deviations (RMSD) of the CAMBR binding region for each configuration from the MD simulations, relative to the extended, alpha-helical pCaN conformation that is compatible with the CaM binding surface. From these simulations, we identify conformations that are amenable for CaM binding (“open” state) and those unsuitable for CaM binding (“closed” state), using a cutoff of pCaN: 7 Å, lpCaN and lpcCaN: 5 Å. We utilized a more restrictive criterion for the longer constructs, as the 7 Å cutoff assumed for pCaN yielded structures that were incompatible with CaM. RMSD values below the cutoff more closely resemble the fully-extended reference structure, whereas values above this cutoff are more compact. As shown in Fig. 3(a-c), all three peptides adopt a small percentage of CaM-compatible configurations as measured by RMSD and the percentages appear to be insensitive to ionic strength. These data additionally indicate that lpCaN (NCPR = 0.088) has the smallest percentage of CaM-campatible structures as assessed by RMSD compared with the bound CaN complex, while lpcCaN (NCPR = 0.264) and pCaN (NCPR = 0.291) have the most.

**Fig. 3.**
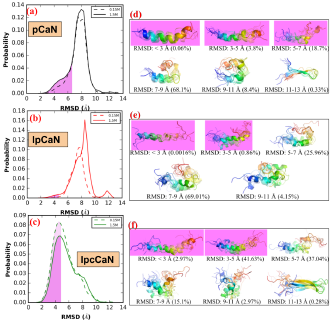
Distribution of RMSD (with respect to bound-pCaN crystal structure in PDB 4Q5U) in the MD of each CaN peptide at 0.15 M ionic strength and 1.5M ionic strength, respectively(a-c). The shaded area colored in violet denotes the open state-like conformation. The representative structures (colored in rainbow with N-termini as blue and C-termini as red) for each RMSD range and percentage of conformations within this RMSD range were also shown(d-f).

In Fig. 3(d-f), we present the structures of the most probable conformations based on RMSD clustering analysis. It is important to note that each peptide configuration was ob-served to partially fold into an *α*-helix indicating that the IDPs considered in this study adopt bound-like ‘residual’ structures in the absence of the binding partner, as has been reported in the literature (29, 43–45). We calculated the fractions of open state conformations sampled in MD simulation based on above mentioned RMSD cutoff (pCaN: 7 Å, lpCaN and lpcCaN: 5 Å), for which we demonstrate that the open states represented a small, but significant fraction (pCaN: ∼20%; lpCaN: ∼1%; lpcCaN: ∼45%) of the conformations sampled. Although the percentage of structures within 3 Å of the bound pose was generally very small (*<* 1%), importantly, these data indicate extended/CaM-compatible conformations are present in the ensemble and that the availability is dependent on the charge density as reported by the NCPR score (see Fig. 3(ac)). We speculate that the tendency for a percentage of the conformation ensemble to assume an extended pose relative to a hairpin fold suggests that intra-molecular repulsion may partially destabilize the formation of loose hairpins. This effect would be exacerbated with charge densities of increasing magnitude, such as those reflected in the NCPR values for pCaN and lpcCaN, and relatively diminished for low NCPR peptides like lpCaN.

To establish a thermodynamic basis for the trends of greater conformational diversity for the high NCPR cases (pCaN and lpcCaN) relative to the low NCPR case (lpCaN), we report potential of mean force (PMF) calculations for these peptides as a function of *α*-helical character, a measure of secondary structure formation, and radius of gyration (R_*g*_), a measure of compactness (see Fig. 4). Such potential of mean force (PMF)s have been used to characterize the propensity for IDPs to assume specific ensemble characteristics, including IDP compactness(46, 47). Given that RMSD distributions shown in Fig. 3 demonstrate nearly identical distributions for 0.15 and 1.5 M ionic strengths, PMF calculations were not performed for the 1.5 M ionic strength. Among the CaN RD constructs considered here, each was observed to present degrees of helical character for the PMF minima that were significantly smaller than those reported for the pCaN peptide when bound to CaM (*α*-helical= 0.84). This suggests that helix formation is thermodynamically disfavored in the absence of CaM. In other words, lacking CaM, unfolded CaN RD states dominate the conformational distribution and thus shifting the conformational ensemble toward the CaM-bound state is thermodynamically unfavorable. This suggests elements of CaM/CaN PPI formation proceed through an induced-fit mechanism.

**Fig. 4.**
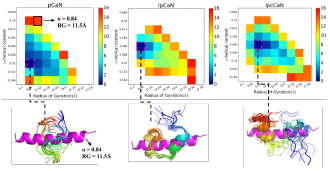
Two dimensional PMFs for pCaN, lpCaN and lpcCaN at 0.15 M ionic strength. The *x* and *y* axes depict *α*-helical and R_*g*_ reaction coordinates, respectively. For each peptide, ten randomly selected structures (colored in rainbow with N-termini as blue and C-termini as red) from lowest energy area are compared against bound state pCaN conformation (colored in magenta) from PDB 4Q5U (*α* = 0.844, R_*g*_ = 11.54 Å). The unit of color bar is *k*_*b*_*T* where *k*_*b*_ is Boltzmann constant and *T* = 298 K is temperature‥

Interestingly, we observe that the range of R_g_and *α*-helical values within a few ***k*_*b*_*T*** of the energy minima (0 ***k*_*b*_*T***) are larger for the high NCPR cases compared to lpCaN. These data mirror our findings for the histogram of RMSD distributions in Fig. 3, with the low NCPR case presenting a narrower distribution relative to the high NCPR cases. Further, the PMF data support the observation for the lpCaN and lpcCaN peptides that the former structure assumes a more compact, hairpin-like configuration relative to the latter, as we observed in Fig. 3(d-f). This indicates that the high NCPR cases access a larger range of conformations in their IDP ensembles that overlap with the CaM-bound structures, albeit in contrast to the more narrowly peaked distributions presented for the low NCPR (lpCaN) configuration. Our results are consistent with the work done by Mao et al(19) for protamine IDPs demonstrating that globule-to-coil transitions were more favored with increasing of NCPR values.

### B. CaN regulatory domain ensemble conformational dynamics are rapid and have modest ionic-strength dependence

Our unconstrained MD and PMF calculations both indicate that the CaN RD peptides do not readily assume an open-state compatible with CaM, although there exist some infrequent, CaM-compatible configurations. In this regard, one can view the accessibility of the pre-folded CAMBR domain to CaM as a ‘gating’ event, which in principle could control the apparent binding rate for this process (48, 49). Given that our previous work (arXiv:1611.04080v1) in which CaN peptides are assumed to have fully CaM-compatible CAMBR conformations demonstrated that all three peptides are capable of binding CaM, albeit at substantially different rates, we postulated here that the kinetics of CAMBR exposure are sufficiently rapid to support diffusion-limited binding. Specifically, we hypothesized that the appearance of bound-like structures before binding serves to nucleate loosely-associated CaM-compatible transient encounter state with low alpha-helical character, which permits “induced folding” in the presence of CaM to access alpha-helix rich bound-states.

As a first step towards probing this hypothesis, we first estimated the transition kinetics between open and closed states identified in Sect. A using Markov state model (MSM). Intuitively, we would expect that higher rates of accessing CaM-compatible open states would maximize the CaM/CaN association rate. We note here that we defined the open state as consisting of conformations below the 7 Å (for pCaN) and 5 Å (for lpCaN and lpcCaN) RMSD cutoff used in Fig. S8 (the red dash lines depict the RMSD of open and closed state for each peptide), while all conformations with dissimilar RMSDs were lumped into a single closed state. We verify that the states are essentially Markovian as the correlation times become negligible beyond roughly tens of nanoseconds (see Fig. S3) which is faster than the diffusion encounter time. Overall, based on this partition of MD data, the transition rates between closed and open states are rapid (the slowest rate is at 1 *×* 10^7^ s^−1^ order, see Table S1) and lead to the short-lived open states shown in Fig. 5 (average life times of open state for all three peptides under both ionic strengths are around 0.2 ns).

**Fig. 5.**
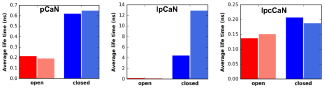
Average life time of open and closed states of CaN peptides determined by MSM at 0.15 M (faded color) and 1.5 M (dark color), respectively. The specific life time and corresponding gating rates are listed in Table S1.

Ionic strength was shown to have negligible impact on the RMSD of our predicted peptide structures relative to CaM-bound conformation. However, given the pronounced role of electrostatics in facilitating protein/protein association rates and protein folding(50, 51), we sought to determine whether transition kinetics between conformations, which highly involve charged species, were influenced by ionic strength. Hence, we compared Markov state model (MSM) rate predictions for MD generated structures at low (0.15 M) and high (1.5 M) ionic strength. Here we found that for pCaN and lpcCaN, increasing ionic strength from 0.15 M to 1.5 M does not affect the gating rates between open and closed states. However, for lpCaN, increasing ionic strength increased *k*f from 1 *×* 10^7^ s^−1^ to 1 *×* 10^8^ s^−1^ order. As a result the lifetime of its closed states decreased from 12.83 to 4.42 ns, as shown in Fig. 5. Hence, the open and closing kinetics of peptides with high NCPR are appear to be less sensitive to ionic strength, compared to structures with low NCPR. These results concur with findings from Liu et al(52), for which they demonstrated that the fast-phase structural fluctuations as measured by Fluorescence correlation spectroscopy (FCR) for the IDP Sic1 disappeared with decreasing ionic strength.

### C. Long-range interactions promote rapid CaM/CaN association

Our results thus far indicate that the CaN RD peptides adopt CaM-compatible conformations in the absence of CaM frequently, albeit transiently. Here we determine the compatibility of these transient states with the CaM/CaN binding interface using Brownian dynamic (BD) simulations. Specifically, we sought to evaluate two hypotheses: 1) that frequent presentation of CaN open states promote near diffusion-limited association rates and 2) that long-range electrostatic interactions are exploited in PPIs involving IDPs. Motivating our first hypothesis are recent indications that target-compatible residual structures of the isolated PUMA IDP form spontaneously as a function of ionic strength and electrolyte composition (53). For the latter hypothesis, we adopt the paradigm of electrostatically-driven association of globular proteins (23, 24, 54, 55), which depends critically on the notion of a transient encounter complex (56, 57). The encounter complex serves as the rate determining step in PPI formation, whereby a protein loosely binds to its protein target, before adopting the fully-formed bound configuration. However, unlike PPIs involving globular partners that typically feature regions of complementarily-charged hydrophilic patches (58, 59), such patches may only be transiently presented, if at all, in IDPs. Nevertheless, ionic-strength dependent binding of IDP-dependent PPIs have been demonstrated (27).

We tested these hypotheses by assuming each peptide must achieve a minimal number of ‘native contacts’ with the CaM N-terminal and C-terminal domains to constitute a transient encounter complex. The native contacts are obtained by analyzing the crystal structure of CaM-pCaN complex, in which key interactions between CaM and pCaN were extracted to guide the BD simulations. From this standpoint, the lenient conditions for association are tantamount to the notion of a transient encounter complex (60, 61), which is formed upon association of two binding partners prior to forming the fully-bound complex. Because we test the first hypothesis using conformations generated from the MD simulations without CaM, this test bears similarity to the conformational selection paradigm (30), though we emphasize CaM is likely required to completely form the bound complex from the transient encounter state. In Fig. S8 and Fig. S7 we show that the MD-generated open states presented in each of the peptide configurations are compatible with both the Nand C-terminal CaM domains to varying degrees, as the open state of each peptide gives BD-simulated kons in the diffusion-limited regime (*>* 1 *×* 10^7^ M^−1^ s^−1^). Notably, all peptides considered here are capable of forming the transient encounter complex with CaM. Further, these rates decrease within increasing ionic strength, as outlined in Sect. C.

We note that modestly high association rates were observed when RMSD *>* 11 Å. We believe these abnormally high rates were artifactual, since visual inspection of these conformations revealed mostly highly compact, bent-coils and *β*-sheets structures (see Fig. 3(d-f) for representative structures in this RMSD range), that might require significant conformational rearrangement to assume the alpha-helix required for CaM binding. However, it is also possible that the structures could help nucleate rapid alpha-helix formation, although we did not explicitly investigate this hypothesis.

Lastly, we investigated the role of conformational gating rates on the effective association rates. Based on the stochastic gating model postulated by Szabo et al (62), there are two limits that bound the effective rates: 1) given gating rates that are significantly faster than diffusional encounter rate (*k*_f_ +*k*_b_≫ *k*_on_), the effective association rate *k*_eff_ is equivalent to the rate associated with the open state, that is, *k*_eff_ = *k*_on_.2) given gating rates significantly smaller than the diffusional encounter rate (*k*_f_ + *k*_b_ ≪ *k*_on_), *k*_eff_ is given by the weighted average of the association rates for all accessed states, that is *k*_eff_ = ⟨*k*_on_⟩. Rates associated with intermediate regimes are obtained by evaluating Eq. 3 using the MSM-estimated gating rates. Based on the data in Table S2 we show in Fig. 6 for pCaN and lpcCaN that *k*_eff_ and *k*_on_ are comparable (e.g. *k*_eff_*/k*_on_ *→* 1), indicating a marginal effect of conformational gating on the association rate. This arises because the conformational transition rates are of the order 1 *×* 10^9^ s^−1^, roughly 100 times faster than diffusional encounter rate, based on our BD simulated *k*_on_s of 1 *×* 10^7^ M^−1^ s^−1^ order (see Table S2). In contrast, the slower transition kinetics for lpCaN yield a *k*_eff_ that is about 50% of the maximal *k*_on_, albeit it is still in a diffusion-limited regime. Moreover, the rates are strongly attenuated at 1.5 M relative to low ionic strength conditions of 0.15 M, which suggest the strong role of long-range electrostatic interactions in promoting association. These data indicate that diffusion-limited association kinetics are realized in the CaN IDP constructs, though the effective rate depends both on ensemble gating kinetics and long-range electrostatic interactions.

**Fig. 6.**
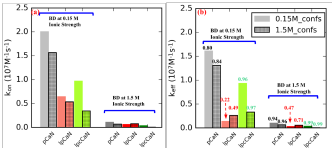
Association rate constants between CaN peptide and CaM before (a) and after (b) taking CaN peptide’s conformational dynamic (from MSM modeling) into consideration using Eq. 3. The bars without grids and with grids depict results in which CaN peptide conformations were sampled at 0.15 M and 1.5 M ionic strength, respectively. In (a), *k*_on_ was calculated via Eq. 2 where *k*_*Cterm*_ and *K*_*Nterm*_ are the average values of 10 randomly selected conformations from each peptide’s open state. In (b) the numbers above each bar represent the ratios of *k*_eff_ to k_on_‥

## 3. Conclusions

Our studies of CaN conformational dynamics and CaM/CaN association reveal several interesting features. While the role of charge distribution in IDPs has been shown to be a strong predictor of ensemble structure including compactness(19, 21), our simulations reveal that measures such as NCPR may offer predictive estimates for the ionic strength sensitivity of conformation transition kinetics. Namely, higher NCPR structures are more likely to adopt conformations that complement their binding target, and are less sensitive to changes in ionic strength that may influence gating kinetics. However, it is important to note that this trend may not generalize to necessarily all IDPs, given the wide range of protein/protein association rates (*<* 1 *×* 10^3^ to *>* 1 *×* 10^9^ M^−1^ s^−1^ (23)) reported in the literature, which hints at the possibility of different assembly mechanisms. Second, we demonstrate that long-range electrostatic interactions can play a paramount role in determining the kinetics of forming PPIs involving intrinsically-disordered partners, while protein-solvent and protein-protein electrostatic interactions govern the kinetics of presenting target-compatible binding motifs. Together, these factors suggest that IDPs can achieve diffusion-limited association by controlling conformational gating, so long as a conformation amenable to association is rapidly sampled. Overall our findings build upon the growing understanding of the roles of conformation selection and induced fit in dictating PPIs, both identifying how conformational selection can accelerate association, despite potential requirements for induced fitting in order to adopt the final binding pose.

Our analyses rest on the core assumption that once the transient encounter complex is formed between the CaN CAMBR and either the N-terminal or C-terminal component of CaM (roughly 100 ns based on the BD simulations), the formation of the final bound complex is comparatively rapid. We justify this from the following: Firstly, while fluorescence correlation spectroscopy (FCS) spectroscopy and molecular dynamics (MD)(63, 64), indicate that the collapse of the extended CaM N/C terminal complexes is believed to occur on the 100 *μ*s timescale in the absence of peptide, we expect that the presence of CaM peptides substantially accelerates the collapse, given that diffusion-limited association of CaM/CaN is reported even at high ionic strengths (arXiv:1611.04080v1). Second, alpha helices, such as those comprising the CAMBR, have been shown to form on the nanosecond timescale based on the coil-to-helix transitions (65) and molecular simulations (66). While nucleation of helix is believe to occur on the millisecond timescale (67), a report has shown that the nucleation and elongation of alpha-helix are on the 20-70 ns and 1 ns timescale, respectively (66). Here, we anticipate that the formation of the transient encounter complex serves as a nucleation event, whereby the initial helical character forms and is followed by rapid helix propagation. Third, though the formation of the bound configuration from the transient encounter complex might be expected to entail substantial resampling and repacking of PPI residues that would slow association, we report in Sect. C folding predictions for pCaN/CaM from a coarse 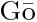 model that demonstrate rapid formation of the fully-folded complex when greater than 50% of the native contacts are formed (see Fig. S9). Here, our poses predicted by BD are ∼55% of *Q*_*n*_ and ∼58% of *Q*_*c*_. Further, upon adopting *Q*_*n*_ and *Q*_*c*_ of this magnitude, the folding landscape predicted in Fig. S9B indicates that the transition toward the fully-folded complex is strictly decreasing in energy. While the kinetics for such a process cannot be exactly determined from the Gō;model, it is believed that minimizing ‘frustration’ along folding reaction coordinates tends to optimize folding rates (68). Finally, experiments summarized in our recent submission(arXiv:1611.04080v1) that CaN peptides bind CaM in a diffusion-limited fashion (2.2 *×* 10^8^ M^−1^ s^−1^ at 0.528 M NaCl) that is accelerated by attractive, long-range electrostatic interactions; even at high ionic strengths for which such interactions are largely attenuated, the association still falls within a diffusion-limited regime (4.39 *×* 10^7^ M^−1^ s^−1^ at 2.028 M NaCl for diffusion-limited regimes (23)). In other words, if the timescale for transitioning from the transient encounter complex toward the bound state were significant, we could expect little sensitivity of the association rate to ionic strength and further, PPI association at high ionic strength would not be expected to be diffusion-limited.

Our study focused on CaN’s binding interaction with CaM, of which the latter regulates a staggering array of eukaryotic signaling cascades through forming PPIs with target protein (5). What sets CaM apart from other such hubs is the surprisingly diverse variety of targets it regulates, despite presenting a single isoform across all mammalian species (69) and seven characterized, disease-associated mutations (70, 71). In part, its ability to regulate this diversity is attributed to the conformational heterogeneity of the CaM binding interface (72) it is capable of forming. Our findings demonstrate that kinetic properties arising from the CaM IDP target independently contribute to the kinetics of PPI formation, and thus constitute an additional tool for tuning the fidelity of CaM-mediated signaling events.

It is important to highlight that formation of the CAMBR/CaM complex alone is insufficient to relieve CaN inhibition by its AID. Namely, Tori et al (11) established that additional interactions between the RD and CaM are necessary to achieve maximal phosphatase activity. While specific interactions between CaM and these RD residues distal to the CAMBR have not yet been determined, it has been suggested that formation of helical character in the RD beyond the CAMBR region correlates with CaN phosphatase activity (11). From this standpoint, CaN activation by way of CaM binding may be triggered by the rapid, diffusion-limited association between the CAMBR region and CaM, which would effectively localize the CaM solvent-exposed surface toward additional RD residues that may extend the PPI interface. In this process, the AID could be ‘reeled in’ after which binding to the CaN catalytic domain would be less probable.

Given the intrinsic disorder reflected throughout the RD domain, it is reasonable to expect that RD ensemble properties and conformation kinetics could tune the final stages of CaM recognition necessary for relieving CaN inhibition. While it is appealing to postulate the phosphorylation or other common post-translational modifications that control charge density might be exploited to modulate the kinetics of this inhibition process, to our knowledge no such post-translational modifications have been identified for CaN. However, we anticipate that proteins that utilize IDP RDs, including the M-domain of myosin binding protein C(73) (residue 262 to 316, NCPR= 0.036), may utilize post-translational modifications including phosphorylation to tune rates of PPI formation.

These findings provide intriguing insight into the interplay between conformational diversity and electrostatically-driven protein-protein association involving CaN, which are likely to extend to subsets of wide-ranging processes regulated by intrinsically-disordered proteins. As such, exploiting IDP composition to tune PPI kinetics could offer new tools to probe and modulate important biochemical signal transduction pathways.

## 4. Methods

### Materials and Methods

#### A. Structure Preparation

The N-domain (residue ID: 3-75) and Cdomain (residue ID: 76-147) of CaN were extracted from the crystal structure (PDB ID: 4Q5U(40)). For CaN peptides, three different peptides with varying lengths and charge distributions were considered: 1) pCaN: native binding region for CaM. 2) lpCaN: elongated pCaN with five additional residues added to two ends of pCaN, respectively. 3) lpcCaN: charge mutated lpCaN having EESE to KKSK mutations at the C-terminal end. Rosetta(74) was used to model initial conformations for the CaN peptides. The *ab initio* structural prediction was conducted by running the ‘AbinitioRelax.linuxgccrelease’ installed on our local host.

The number of output conformations was set to ten. No predefined secondary structure file was specified. The rest parameters were not explicitly specified (namely using their default values, as that listed in (75)). According to the energy score, for each CaN peptide, the conformation with lowest energy was picked out for further extensive MD sampling. Although just one conformations was selected for each CaN peptide, it was expected that the following microsecond MD ensure adequate sampling.

#### B. Molecular Dynamic Simulation

We next performed MD simulations to extensively explore the conformational spaces of the CaN peptides. The Amber ff99SB-ILDN(76) force field which improves the amino acid side chain potential in IDP was used in the simulations. Due to the intrinsically disordered nature of these peptides, special care should be taken when choosing force fields since current force fields would sample conformations that are over collapsed when applied on IDPs(77). The MD was performed by using Amber14(78). The implicit solvent model (igb = 2 with salt concentration = 0.15 M) was used. The cutoff value for non-bond interactions was set as 999 Å. The starting structure was first subjected to 50000 steps of energy minimization. The minimized structure was slowly heated from 1 to 298.15 K by using the Berendsen Thermostat within 800 ps. During the MD process, the time interval was set to 2 fs and the SHAKE(79) constraints were applied on bonds involving hydrogen atoms. By setting the initial temperature in the heating stage equal to 1 instead of 0 and ig = −1 would generate different initial velocity distributions for the system, thus independent simulations can be achieved. For each peptide, three independent MDs were performed to ensure the reliability of the sampling (total 15 *μ*s production run for each peptide). To study the effect of ionic strength on sampling, we ran analogous simulation with salt concentration = 1.5 M, resulting in a total 30 *μ*s production run for each peptide (15 *μ*s at 0.15 M and 15 *μ*s at 1.5 M ionic strength).

#### C. Two-dimensional replica-exchange umbrella sampling (REUS) PMF calculation

Two-dimensional PMF calculations were performed to characterize the free energy profile associated with conformational space of each peptide. Two reaction coordinate (RC)s were defined: 1) *α* which describes the *α*-helical content of the peptide (ranging from 0.1 to 0.9) and 2) R_*g*_ of the peptide (ranging from 5 to 32 Å). Each RC range was divided into nine bins resulting in total 81 windows (with interval being 0.1 and 3Å for *α* and R_*g*_, respectively). The two force constants of the harmonic potentials imposed on these two RCs are 1.000 *×* 10^3^ kcal mol^−1^ U^−2^ for *α* and 2.5 kcal mol^−1^ Å^−2^ for R_*g*_. For each peptide, the representative structure from the most populated cluster was chosen as the starting structure. NAMD2.11(80) was chosen to perform the 2D replica-exchange umbrella sampling (REUS) calculations due to it’s *colvar* module which supports various user-defined collective variables. The CHARMM36(81, 82) force field was used in the 2D REUS calculations. For each window, the simulation length was set to 20 ns and only the last 15 ns data was used to calculate free energy by WHAM(83).

#### D. Markov state model

(MSM) analysis via Aqualab. For each peptide, a 1D kinetic trajectory was created from the 15 *μ*s MD trajectory describing the state change along simulation time. For pCaN, the open state was defined as RMSD *<* 7 Å while for lpCaN and lpcCaN the open state are defined as RMSD *<* 5 Å (this state definition criterion was later justified by Fig. S8 and Fig. S7 in which open state conformations have obvious larger simulated *k*_on_ for both with CaM N and C domains). A kinetic network was created based on the 1D kinetic trajectory via Aqualab (84), which seeks to impose the detailed balance condition *P*_*i*_*T*_*ij*_ = *P*_*j*_ *T*_*ji*_, where ***P*** is equilibrium probability matrix and ***T*** is transition probability matrix.

#### E. Brownian dynamic

(BD) Simulations. The binding of CaN peptide and N/C terminal domains of CaM are treated as two independent events and simulated separately by using the BrownDye package(85). For each peptide, ten conformations for each RMSD cluster were randomly selected to perform BD simulations with N/C-domain of CaM. The PDB2PQR(86) was first used to generate the pqr files for CaM N/C domains and the selected conformations of CaN peptides from MD trajectory with radii and point charge parameters adapted from the AMBER99(87) force field. The generated pqr files were then passed into APBS(88) to evaluate the electrostatic potential of these structures. APBS was used to numerically solve the linearized Poisson-Boltzmann equation assuming an ionic strength of 0.15 M and 1.5 M NaCl:

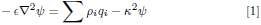

where *ψ* is the electrostatic potential, *ρ*_*i*_*q*_*i*_ is the charge distribution of fixed charge *i*, and *κ* is the inverse of Debye length. The Debye length reflects the scale over which mobile charges could screen out electric potential fields.

In present BD simulation, the reaction criterion was chosen to be six pairs of contacts with distance of contact being less than 10 Å. The contact list was created via the make_rxn_pairs routine in Browndye package based on the pCaN-CaM complex crystal structure (PDB ID: 4Q5U) with distance cutoff being 5 Å. 10000 single trajectory simulations for each system were conducted on 10 parallel processors using nam-simulation. Thus for each peptide, the total number of BD trajectories was about 1 million. The reaction rate constants were calculated using compute-rate-constant from the BrownDye package.

To estimate the association rate and its sensitivity to ionic strength, we computed association rates for the terminal domains separately, assuming that both components bind independently,

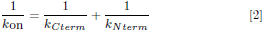

where the rates in the right hand side correspond to the association rates for the C and N terminal domains, respectively. We anticipate that this expression under-approximates the rate of complex formation, given that tethered binding partners generally exhibit higher efficiencies for forming intact complexes (89, 90).

#### F. Effective Association Rate Combined with Gating Kinetics

The effective association rate constant after taking conformational dynamics into account was given by Szabo et al(62):

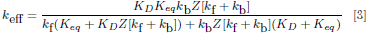

Where

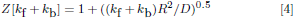

where *K*_*D*_ is the association rate when the peptide is always in open state and in present study, *K*_*D*_ is the BD simulated association rate constant of the open state CaN peptides with CaM (e.g., *K*_*D*_ = *k*_*on,open*_). *K*_*eq*_ is characteristic constant indicating the extent to which the association is diffusion-controlled (see (62) for more details). In present study, we set *Keq* = 1 *×* 10^20^ M^−1^ s^−1^ and justified by showing in Fig. S6 the sensitivity of *k*_eff_ to *K*_*eq*_, as we can see that *k*_eff_ values for all peptides become fixed after *K*_*eq*_ is larger than 1 *×* 106 M^−1^ s^−1^. *k*_f_ and *k*_b_ are the conversion rate between the open and closed state determined from MSM. *R* is the contact distance at which the transient complex formed and in present study we set *R* equal to the average b-radius values from BD simulations. *D* is the relative translational diffusional constant and was calculated via (23, 85).

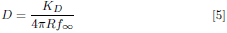

where *f*_*∞*_ is the reaction probability which was at the order of 1 *×* 10^−4^ given by BD simulations.

## ACKNOWLEDGMENTS.

This article is dedicated to the memory of late Professor Jeffry A. Madura. Research reported in this publication was supported by the Maximizing Investigators’ Research Award (MIRA) (R35) from the National Institute of General Medical Sciences (NIGMS) of the National Institutes of Health (NIH) under grant number R35GM124977. This work used the Extreme Science and Engineering Discovery Environment (XSEDE)(91), which is supported by National Science Foundation grant number ACI1548562

## 5. Supplementary Information (SI)

### A. Supplementary figures

**Fig. S1.**
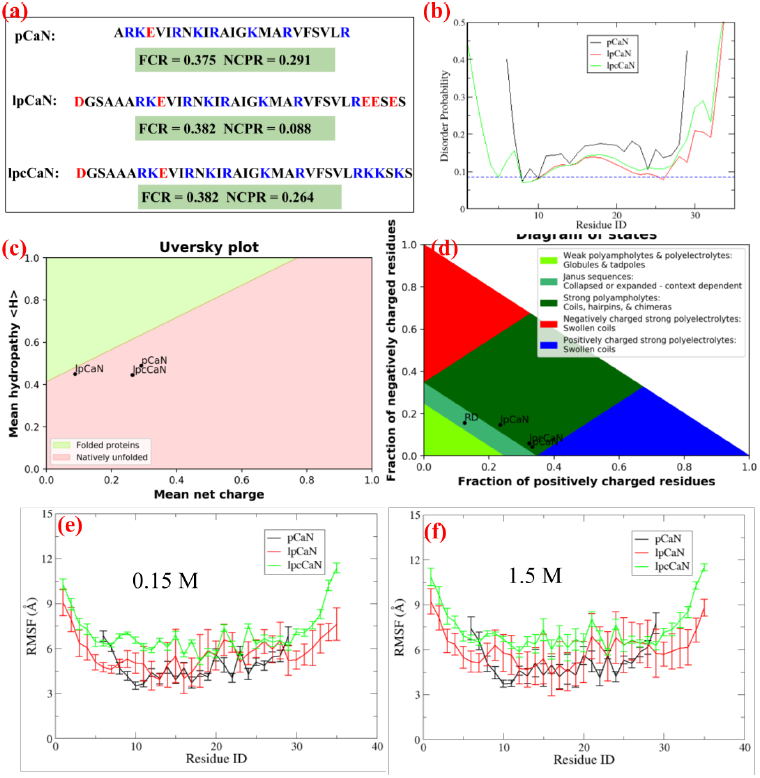
(a) Sequences of the three CaN peptides studied in the present work. The positively charged residues and negatively charged residues are colored in blue and red, respectively, along with FCR and NCPR scores. (b) Disorder probabilities predicted by DisEMBL(41). The shown curves are scores based on “hot-loops” which is reported to be a good criterion to define disorder(41, 92). The blue dash line denotes random expectation values. (c) Mean hydropathy score and mean net charges of the three peptides and their locations in the Uversky diagram(93) (d) Locations of the three CaN peptides and CaN RD in the Das-Pappu diagram(94). Figures (c) and (d) are generated by localCIDER(42). (e-f) RMSF of each residue during the 15 *μ*s MD at 0.15M and 1.5M ionic strength, respectively.

**Fig. S2.**
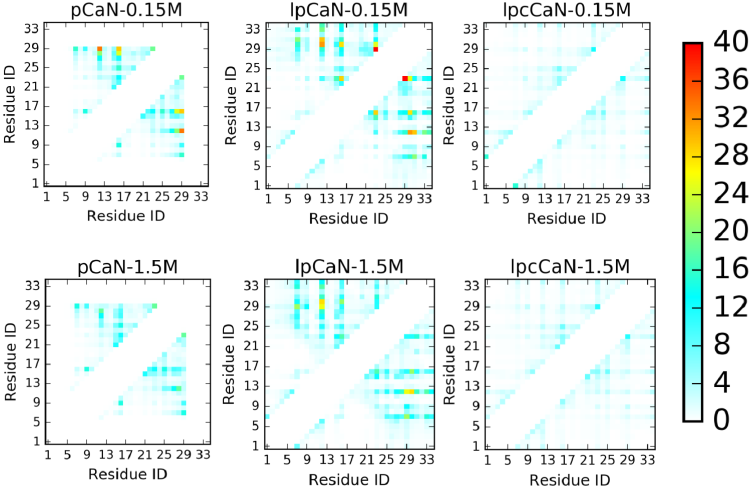
Contact map analysis of 15 *μ*s MD trajectory of three CaN peptide under 0.15 M and 1.5 M ionic strength, respectively. Contact data was collected via CPPTRAJ in Amber with distance cutoff as 7 Å and only residue pairs which are at least 5 residues apart (*i* and *i* + 5) in sequence are considered. The unit of numbers on color bar is number of average contacts for each pair over the simulation time.

**Fig. S3.**
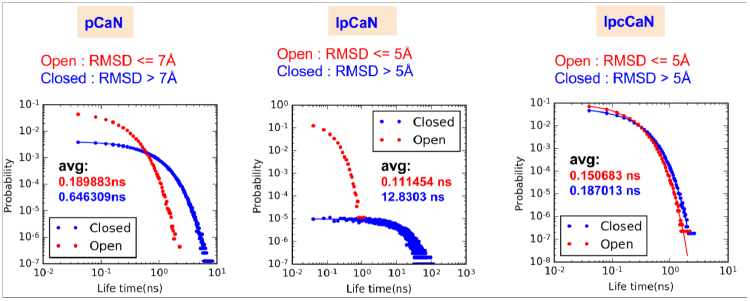
Life time distribution of open and closed states for each CaN peptide calculated from the 15 *μ*s MD trajectory (0.15 M ionic strength) by Aqualab using the MSM model‥

**Fig. S4.**
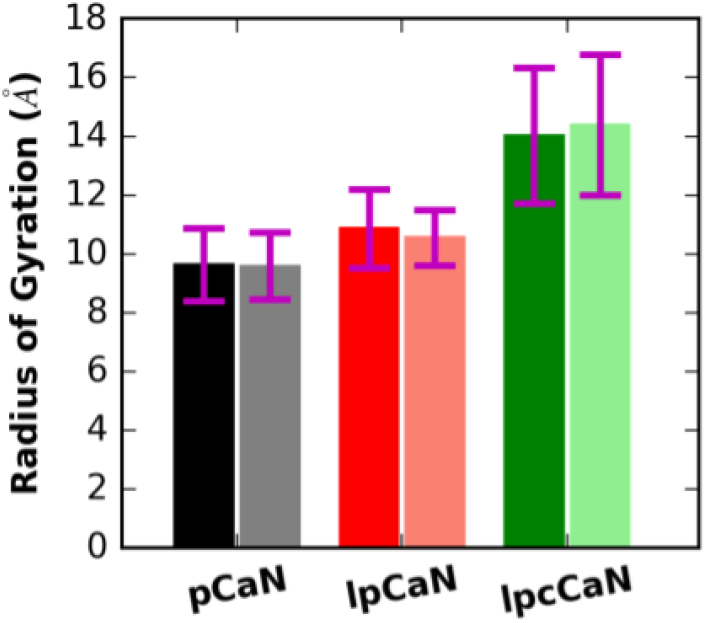
Average R^*g*^ of CaN peptides at 0.15 M (faded color) and 1.5 M (dark color), respectively‥

**Fig. S5.**
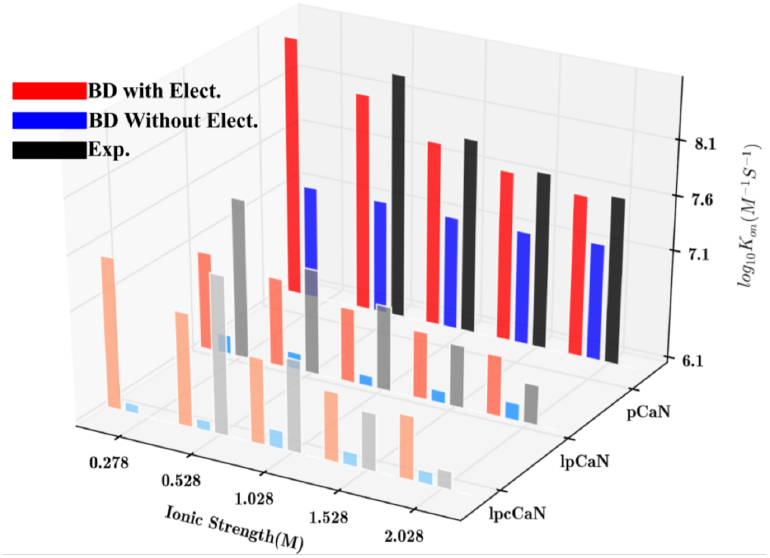
B.calculated association kinetics between rigid CaN peptides and CaM. The electrostatic interaction was turned off by setting CaN peptides charges to be zero. Specifically, after turning of electrostatic interaction, lpCaN retains 50% above association rates while lpCaN and lpcCaN reduce to much smaller *k*_on_s, implying that electrostatic interaction has larger impacts on lpCaN and lpcCaN than pCaN‥

**Fig. S6.**
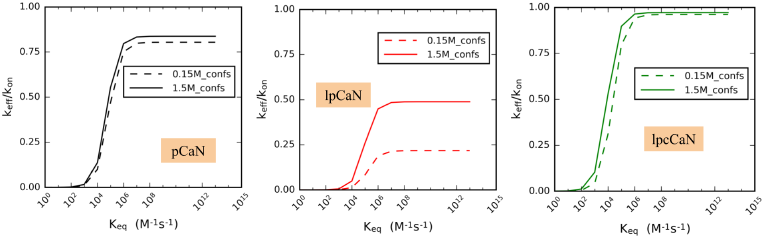
Sensitivity of *k*^*eff*^ to *K*^*eq*^ for three CaN peptides sampled at 0.15 M and 1.5 M ionic strength, respectively‥

**Fig. S7.**
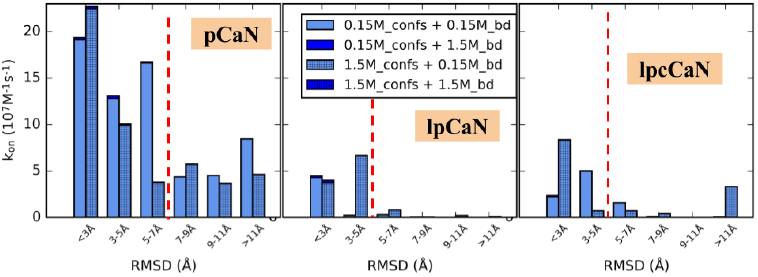
BD-simulated separate association rate constants with CaM C domain vs. RMSD under 0.15 M and 1.5 M ionic strength with CaN peptides conformations sampled at the same two ionic strengths, respectively. The red dash line designates the border of open and closed states based on RMSD‥

**Fig. S8.**
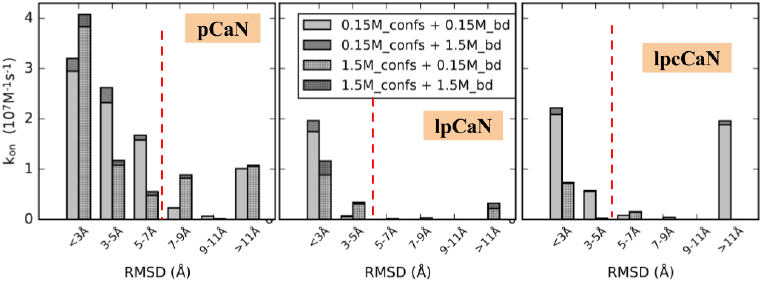
BD-simulated separate association rate constants with CaM N domain vs. RMSD under 0.15 M and 1.5 M ionic strength with CaN peptides conformations sampled at the same two ionic strengths, respectively. The red dash line designates the border of open and closed states based on RMSD.

Given that all-atom simulations of CaM-induced folding of CaN to form the fully-bound complex are intractable, we utilized a Gōmodel to infer the relative compatibility of the BD-generated transient encounter poses with the final binding pose. Here the CaM-pCaN crystal complex structure (PDB ID: 4Q5U) was used to define native contacts and topology based on the standard protocol for a Gō modal adopted by Karanicolas-Brooks(95). As shown in Fig. S9A, for the coarse-grained (CG) simulations, *Q*^*n*^ first becomes larger than zero, which suggests that pCaN first partially interacts with N-domain of CaM before interacting with the C-domain. Additionally Fig. S9A shows that the fully-bound pCaN/CaM complex forms at the timescale of ∼50 ns, which is comparable with the ∼100 ns diffusional timescale obtained by BD simulations. Importantly, we demonstrate in Fig. S9A that the BD generated transient complexes already exhibit *Q*_*n*_ and *Q*_*c*_ around 55%, which are sufficiently high to rapidly access the fully-folded complex. This is supported the 2D free energy profile in Fig. S9B computing using the Gō model, for which paths originating with *Q*_*n*_ and *Q*_*c*_ values above roughly 50% strictly decrease toward the final bound state.

**Fig. S9.**
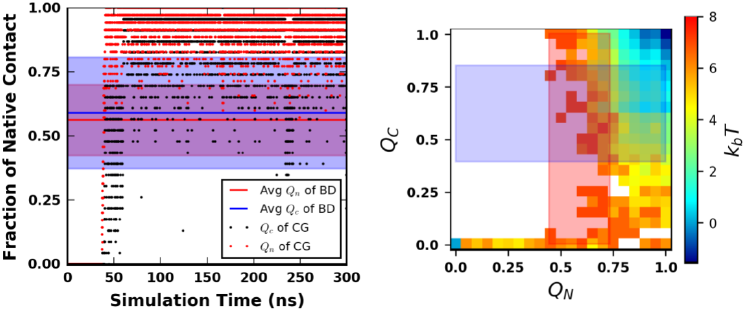
Fraction of native contact (*Q*) and free energy profile in CG simulation with Gō model. A: Fractions of native contact between N/C-domain of CaM and pCaN (denoted as *Q*^*n*^ and *Q*^*c*^, respectively) along CG simulation time (only the first 300 ns of 1 *μ*s is shown for clarity). B: 2D free energy profile projected along *Q*_*n*_ and *Q*_*c*_ in CG simulation. The shaded areas colored in light red and blue depict the ranges of *Q*_*n*_ and *Q*_*c*_ values in BD simulations, respectively. The last frame of each BD trajectory of the 10 pCaN open conformations (RMSD *<* 3 Å) with CaM N/C domain were used to calculate average and standard deviation of *Q*_*n*_ and *Q*_*c*_.

### B. Supplemental tables

**Table S1.** Average life times and gating rates between CaN peptides’ open and closed conformations sampled at 0.15 M and 1.5 M ionic strength (sate definition is based on RMSD see red dash lines in Fig. S8 and Fig. S7).

| | Ionic Strength(M) | Average Lifetime (ns) | | Gating rate ($s^{-1}$) | |
| --- | --- | --- | --- | --- | --- |
| | | $\Gamma_{open}$ | $\Gamma_{closed}$ | $k_b$ | $k_f$ |
| pCaN | 0.15 | 0.19 | 0.65 | $5.27 \times 10^9$ | $1.55 \times 10^9$ |
| | 1.5 | 0.21 | 0.62 | $4.70 \times 10^9$ | $1.62 \times 10^9$ |
| lpCaN | 0.15 | 0.11 | 12.83 | $8.97 \times 10^9$ | $7.79 \times 10^7$ |
| | 1.5 | 0.13 | 4.42 | $7.48 \times 10^9$ | $2.26 \times 10^8$ |
| lpcCaN | 0.15 | 0.15 | 0.19 | $6.64 \times 10^9$ | $5.35 \times 10^9$ |
| | 1.5 | 0.14 | 0.21 | $7.31 \times 10^9$ | $4.85 \times 10^9$ |

**Table S2.** BD-simulated encounter rates (*k*on) under 0.15 M and 1.5 M ionic strength for the open state CaN peptides sampled the same two ionic strengths. The effective association rates (*k*^*eff*^) calculated via Eq. 3 are also shown.

| | | $k_{on}/k_{eff} (1 \times 10^7 M^{-1} s^{-1})$ | | Gating rate ($1 \times 10^7 s^{-1}$) | |
| --- | --- | --- | --- | --- | --- |
| | | 0.15_BD | 1.5_BD | $k_b$ | $k_f$ |
| pCaN | 0.15_confs | 2.00 /1.61 | 0.11/0.102 | 527 | 155 |
|  | 1.5_confs | 1.56/1.31 | 0.07/0.065 | 470 | 162 |
| lpCaN | 0.15_confs | 0.64/0.14 | 0.06/0.03 | 897 | 7.79 |
|  | 1.5_confs | 0.53/0.26 | 0.08/0.06 | 748 | 22.6 |
| lpcCaN | 0.15_confs | 0.97/0.93 | 0.04/0.04 | 664 | 535 |
|  | 1.5_confs | 0.34/0.33 | 0.005/0.005 | 731 | 485 |

### C. Supplementary results

Given the enrichments of negatively charged residues in CaM and positively charged residues in CAMBR of CaN, we postulated that long-range electrostatic interaction between these two binding partners should play an important role in association kinetics. Here we considered four configurations from combining BD simulations at 0.15 M and 1.5 M ionic strength (two cases) and MD-generated conformations at the same ionic strengths (two cases). This approach permits the evaluation of our hypothesis that long-range electrostatic accelerate protein-protein association for the CaN IDP constructs, while ionic strength controls the gating of CaN peptide ensembles. As shown in Fig. S8 and Fig. S7, for each conformation, BD simulated *k*_on_ at 0.15 M ionic strength are much larger than analogous results at 1.5 M ionic strength, demonstrating that long-range electrostatic interactions are the predominant driving force behind CaM and CaN association. In contrast, results obtained using high (1.5 M) ionic strength are much slower than these at 0.15 M (see Fig. S8 dark bars). We confirmed these results in Fig. S5 for which we disabled electrostatic contributions, after which the predicted association rates were minimized. Overall, these data implicate long-range electrostatic interactions in driving rapid, diffusion limited protein/IDP association, as is well established for their globular counterparts.(55).

